# Understanding Virtual Staining with generative adversarial networks for Osteoclast Imaging

**DOI:** 10.1101/2025.10.06.679965

**Authors:** Katharina Schmidt, Antonia Obersteiner, Max von Witzleben, Michael Gelinsky, Juergen Czarske, Nektarios Koukourakis

## Abstract

Virtual staining with generative adversarial networks is an efficient, non-invasive and scalable alternative to conventional cell staining, minimizing the need for destructive and time-consuming protocols. In this study, we investigate the explainability of a network trained to virtually stain osteoclast cultures, using intensity-based label-free input images. The model enables analysis of cell cultures without immunostaining. Explainability assessments, including receptive field and feature map analyses, show that the background in input images significantly influences staining predictions within cellular regions and the trained network performs an internal segmentation during the image transformation process. This suggests that contextual cues beyond cell boundaries are implicitly learned and integrated during training. By eliminating repetitive staining procedures, virtual staining enables longitudinal studies, allows multiplexing of individual samples, and reduces reagents and laboratory waste. Our findings enhance understanding of the virtual staining process and highlight its potential for biomedical research applications.

## Introduction

Immunostaining of biological samples is a fundamental tool in biomedical research and clinical diagnostics, enabling the spatial localization of specific proteins within tissue and cells. Despite their relevance for cellular phenotyping, respective staining protocols are labor-intensive, time-consuming, and require costly reagents and extensive manual optimization^1^. Moreover, variability in staining quality and interpretation can limit reproducibility across laboratories^2^. These limitations become even more pronounced in the context of advanced tissue models and regenerative implants, which are increasingly used to study dynamic processes such as differentiation, remodeling, and cell–material interactions under physiologically relevant conditions.

Traditional staining protocols, which aim to visualize such processes usually rely on either chemical or genetic methods (see fig. 1). Chemical stainings are often endpoints of the samples, not allowing longterm studies due to toxicity and thereby induced cell death. In contrast, samples with genetic modification can often be imaged alive, but require extensive resources. In addition, the destructive nature of fixation and labelling steps restricts further sample use in long-term studies especially when working with scarce number of tissue-derived (primary) cells, as these processes irreversibly alter or consume the sample, precluding repeated analysis. Also, the application of multiple stains in one sample increases the number of required cultures and materials overall. Furthermore, multiple stains are in some cases not possible due to cross staining, autofluorescence, chemical interactions, or physical fragility of the sample^3^.

**Figure 1:**
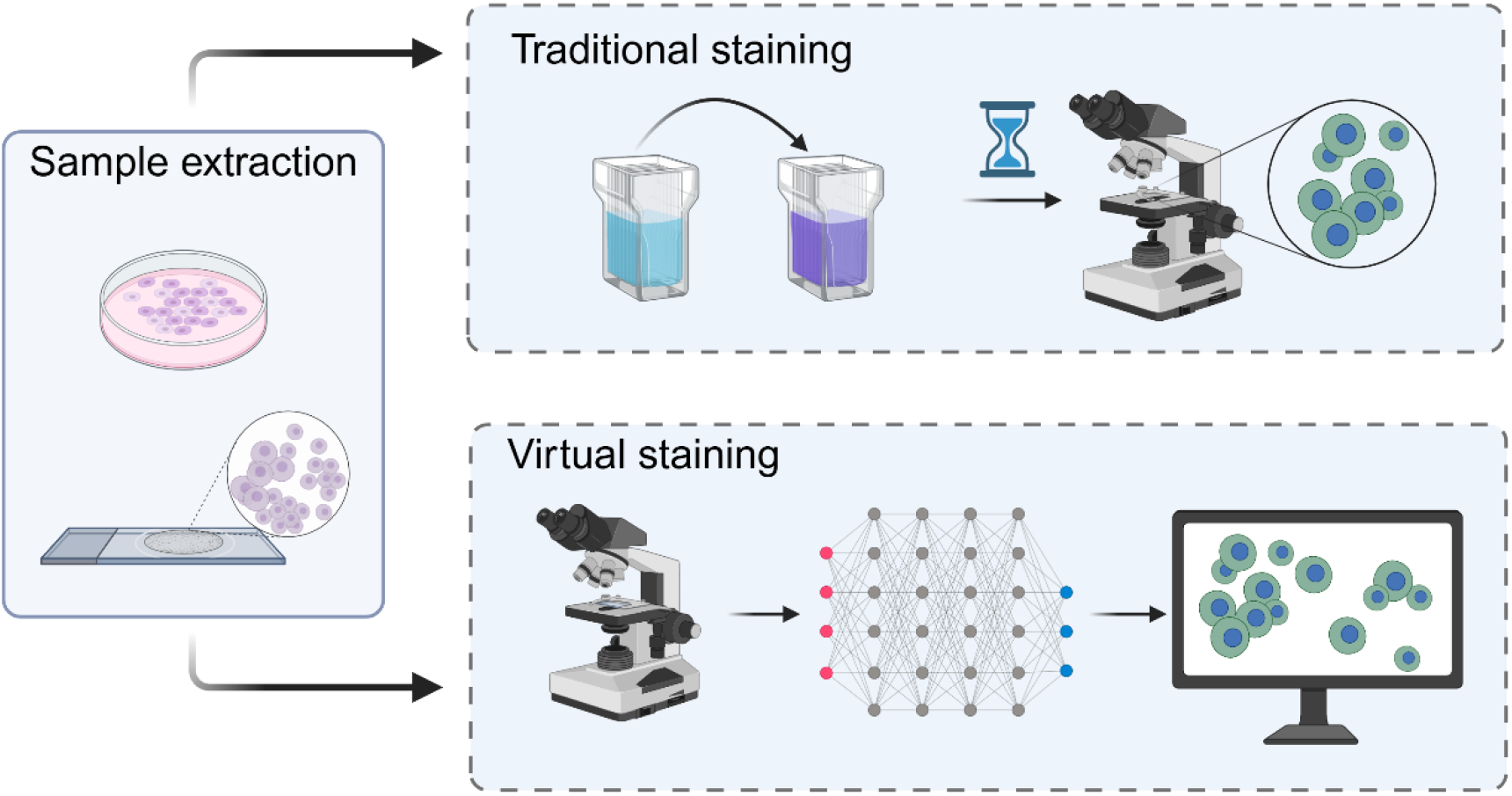
Principle of virtual staining to save time and resources compared to traditional staining protocols. In the upper row staining following a traditional protocol including different staining agents and washing solutions is shown. The imaging is conducted, after the protocol is finished. This slows down the whole process and requires plenty of resources. In comparison, the proposed virtual staining approach is illustrated in the lower row. The imaging is performed on the unstained sample and the captured image is transformed by a neural network to obtain the stained image.

The underlying chemical reactions of traditional staining are highly non-linear processes, therefore neural networks as a form of deep learning are a suitable solution to mimic this. One strategy of digitally staining label-free images is to perform a deep learning assisted segmentation followed by colouring of the segmented areas other methods employ direct image transformation^4^. Such deep learning models—most notably neural networks and generative adversarial networks (GANs)^5^—are capable of learning mappings between label-free microscopy images and their virtually stained counterparts, without the need of chemical or genetical modification^6-12^. In a GAN architecture two networks called generator and discriminator networks are trained against each other^10,13-15^ (see fig. 2a)). The generator predicts an output (fake) image based on given labels or input images. The discriminator needs to decide afterwards if an image is real (from ground truth data) or fake (produced by the generator). The success-rate of the discriminator decision is then included in the loss calculation of the generator network. So, the aim of the generator network is to produce images, which the discriminator cannot tell apart from the ground truth images. This training process enhances the generator’s ability to produce realistic outputs that closely resemble ground truth data. Using these techniques enables digital transformation of images to mimic different stains, which were presented to the network during training. Here, GANs are superior to deep neural network in terms of realistic image predictions, due to the adversarial training process inherent in their framework^10^.

**Figure 2:**
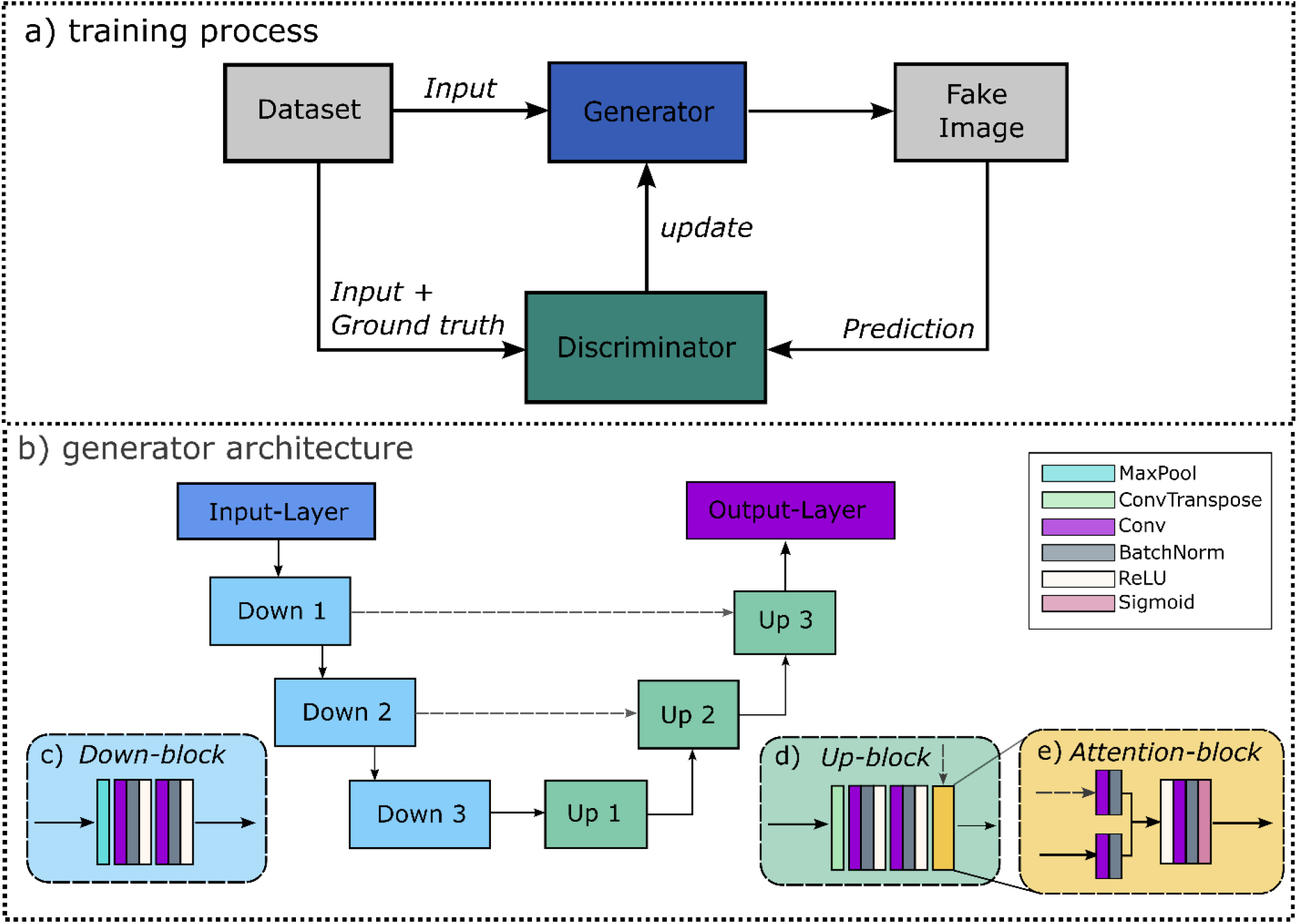
a) Overview of the training process for a generative adversarial network (GAN) with two neural networks called Generator and Discriminator, which are trained against each other. b) architecture of the Pix2Pix Generator network including downsampling blocks, upsampling blocks and skip connections. c)-d) Detailed layer architecture of the used blocks.

In recent studies, neural networks were trained to transform optically sectioned images of unlabelled cells to haematoxylin and eosin (H&E) stained images^11^ or auto-fluorescence images to H&E^6,7,12^. Another significant use-case of virtual staining involves stain-to-stain transfer, where images are transformed from one staining modality to another^6,16-19^. Despite growing interest, current research in virtual staining has focused predominantly on histological applications — especially the reproduction of H&E staining in tissue sections ^6,7,20-22^. These data offer structural consistency and well-reproducible features with minimal variation in staining quality, making them well suited for model training. In contrast, virtual immunostaining in fluorescence-based immunocytochemistry workflows has already shown promising results, but remains far less explored than in histological applications. While histological staining primarily reveals morphological structure, immunohistochemistry and immunofluorescence enable molecular specificity by targeting proteins or subcellular components with labelled antibodies or chemical probes. In the present work, we employed fluorescence-based chemical stains, specifically phalloidin-488 for actin cytoskeleton and 4’,6-Diamidin-2-phenylindole (DAPI)^23^ for nuclear DNA. Further examples for fluorescent modifications are green fluorescent protein and mCherry expressing cells^24-26^. For specific markers of breast cancer such as HER2 and Ki-67 recent publications demonstrate the successful application of virtual staining techniques in immunostaining^27,28^.

In biomedical applications, the trustworthiness of the network’s predictions is a fundamental requirement. However, neural networks are often referred to as black boxes^29^, because it’s unknown how and why a certain output is produced. Despite recent publications about explainability of neural networks in general^30^, the application of these methods to trained virtual staining networks and corresponding conclusions are still missing. In this study, we investigate whether computational models can substitute for conventional nuclei (DAPI) and cytoskeletal (Phalloidin) staining in tissue engineering using a virtual staining approach. To establish a well-controlled baseline, we focus on osteoclasts cultured on standard tissue culture plastic without biomaterials, where staining is technically straightforward and well-characterized. We aim to assess whether a deep learning model in form of a GAN can accurately infer biologically relevant staining patterns and to uncover the underlying key decision-making principles. This study represents a foundational step toward extending virtual staining to more complex, variable, and biomedical research relevant cell-material systems.

## Results

In our study, a GAN architecture was trained for virtual staining of fluorescent cell samples. On the test dataset, which was not shown during training, the trained generator network achieved an average Structural Similarity Index Measure (SSIM) of 0.83 and a peak-signal-to-noise Ratio (PSNR) of 22.475 db. Fig. 3a)-f) illustrates representative predictions alongside ground truth images and input data. Quantitatively, most of the cell structures are recognised and stained correctly, indicating a high concurrence between ground truth and prediction. However, hallucinations such as shifted cell nuclei occur (see red markers in fig. 3b) and 3c) and Figure 1 in the supplement). In fig. 3d)-f), the nucleus and the cytoskeleton are completely detected and coloured by the network, which is the case for most of the test images as indicated by the high SSIM. Detailed views of the hallucinations present in fig. 3a)-f) are provided in fig.2 of the supplement.

**Figure 3:**
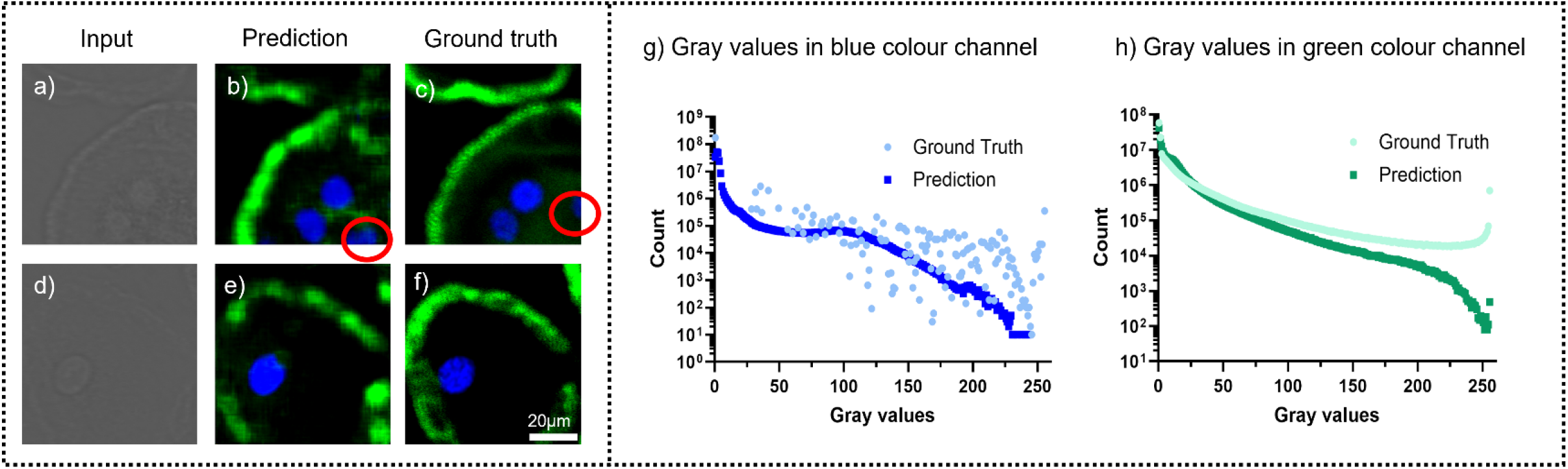
a)-f) examples from the test data, supporting qualitatively high correlation between network prediction and ground truth images of the test data (test SSIM = 0.83). a) and d) are labelfree images captured with a confocal microscope, b) and e) are the predictions of the trained network, c) and f) are fluorescent images with DAPI and Phalloidin staining captured with the same confocal microscope as the input images. g) and h) display the distribution of intensity values for the blue and green colour channel for ground truth and predictions of the test set. During training the network did learn to mimic the intensity distribution of the green colour channel (see h)) and copies this pattern for the blue colour channel (see g)). Note that the ground truth values of the blue colour channel show larger variance compared to the ground truth data from green colour channel.

Furthermore, the colour intensities of the blue and green channel between ground truth and predicted images were compared (see fig. 3 g)-h)). The predicted pixel intensities for the green channels align highly with the ground truth values. In contrast, the ground truth values of the blue channel are more scattered and where not reproduced by the network. The network replicates the distribution of intensities learned by the green colour channel also for the blue color channel. The observed discontinuities may result from the staining characteristics of the respective dyes. In general, DAPI exhibits an intense staining pattern due to its favorable diffusion behavior and strong binding to DNA, typically resulting in higher signal intensities in the blue channel. In contrast, although Phalloidin also provides high staining quality, but manifests as stronger signal at the cell periphery and weaker signal toward the center, leading to less counts of intense signal in the green channel.

For further investigating the functionality of the network and to shed light into which information leads to which staining we tested different augmentations mimicking different cell characteristics.

### Augmentations

The predictions of a trained network depend on the input given to it. So relevant visual features determine the network’s prediction. To distinguish these relevant features from irrelevant ones, studies on augmented input images were conducted. These show in particular the sensitivity of the network to certain input features and also reveal the networks selectivity. The aim of this experiment is to determine the morphology of recognized input features. This information can be used to optimize the image acquisition process in the future and thereby improve the input images for virtual staining.

In the input images the cell nuclei are visible as ring-like structures, as evidenced in fig. 3 a) and d). Based on this finding, a parameter study with different properties of artificially inserted rings was performed. The trained network’s response to variations such as radius, intensity, and border width, is summarized in fig. 4 a)-c), examples of augmentations and corresponding predictions are provided in the methods section. Rings with a radius of 20 pixels, high intensity (grey value > +8% of the average image intensity) and a border width of 7 pixels are coloured most intensely by the network.

**Figure 4:**
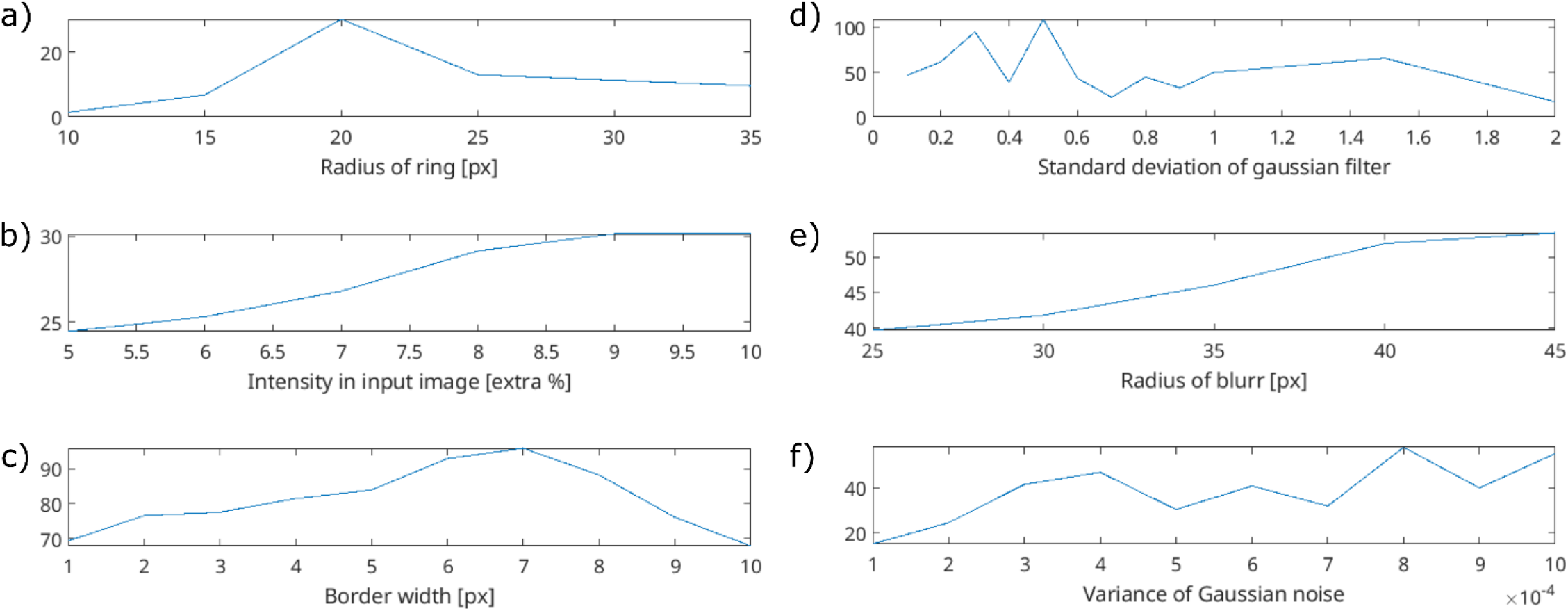
Mean intensities of predictions in the blue (a)-c)) and green (d)-f)) colour channel based on augmentations. In a)-c) rings of varying size, intensity and border width were added. In d)-f) circles with varying size and noise type were added. The y-axis shows the average intensity value of the investigated colour channel for the augmented areas in the predictions.

In the input images, such as the ones in fig. 3, the cytoskeleton is visible as a large spatial distribution of high and low intensity values. Therefore, an additional noise term, which exhibits exactly these characteristics, was used as augmentation to mimic the impression of cytoskeleton and added to a certain area in the input image. These areas in the input images were successfully coloured green by the network. The dependence of noise type and area size are presented in fig. 4d)-f). In this study, Gaussian and Poisson-noise were investigated, to check whether the type of noise has an influence on the network prediction. Furthermore, the influence of the intensity values was tested by variation of the variance for Gaussian noise itself and the standard deviation of a gaussian filter added to the Poisson noise. Fig. 4 d) displays the outcome of regions modified with Poisson noise, which was additionally blurred using a Gaussian filter with a certain kernel standard deviation as tuning parameter. Low standard deviations of the filter lead to high spatial frequencies in the intensity values of the augmented areas. This does quantitatively not align with the noise of the cytoskeleton in the input images and did result in random intensity values in the prediction. From a standard deviation of 0.7 on a clear trend towards a peak at 1.5 is visible. Beyond this the predicted colour intensity decreases, so the trained network does not recognise the intensity distribution as cytoskeleton anymore. This suggests that the predictions of the network highly depend on the spatial frequency distribution of the intensity values representing the cytoskeleton. So, the cell organelles as well as cytoplasm have an influence on the network’s staining decision. This emphasizes the dependence of the staining quality on the representation of cell types in the training dataset used.

In fig. 4 e) the mean intensity for circles with Poisson noise and increasing radius is shown. The mean staining intensity is clearly increasing with the radius, which indicates a stronger recognition of larger circles as cytoskeleton. Fig. 4 f) presents the results for circles with Gaussian noise and a constant radius of 40 pixels, here the network response to the variance of Gaussian noise was investigated. According to the absolute values of mean intensities achieved for the two different kinds of noise (see y-axis of plots 4 d) – f)), the network predicted Poisson noise regions with Gaussian filtering more intensely than those with Gaussian noise alone.

### Receptive fields

The theoretical receptive field (TRF) describes the maximum possible spatial extent in the input microscopy image that could, in theory, affect the network’s staining decision for a particular output pixel. The TRF of the generator network (for the architecture see fig. 2 b)) is governed by the interplay between the kernel sizes in the convolutional layers and the total number of layers, i.e., the network depth^31^. Calculations result in an TRF size of 96 pixels, which is a surface area of 96^2^ = 9216 *px*. Details of the calculation are provided in the supplement.

In contrast, the actual receptive field (ARF) refers to the effective region in the input image that has a non-negligible influence on the output pixel’s prediction^32^. The ARFs of the generator network were determined using guided backpropagation (GBP), details on this are provided in the supplement. Fig. 5a)-f) shows examples from the test dataset, where a single pixel of the output image was backpropagated. In the GBP images (fig. 5 d)-f)) the gradients from the input image which influenced the staining decision for the pixels marked in the prediction image are visualized, the grey values indicate the corresponding gradient value. This experiment reveals that the entire nucleus structure is crucial for the network’s staining prediction of a single pixel inside the nucleus (see fig. 5 f)). In contrast, the staining decision for green-labeled cytoskeletal pixel does not rely on the entire cellular structure, but is predominantly driven by directly adjacent pixels and local features in the immediate neighbourhood (see fig. 5 d) and e)). The correct (fig. 5 d)) and incorrect (fig. 5 e)) prediction of the cytoskeleton have a similarly large receptive field size and do not show a specific pattern such as the roundish structure in fig. 5 f) for the nucleus. The images 5 d) and g) display a shape with two spikes and a central point for correctly estimated cytoskeleton. Figure 5 e) and h) meanwhile show an irregular, lengthy shape for the not recognized part of the cytoskeleton.

**Figure 5:**
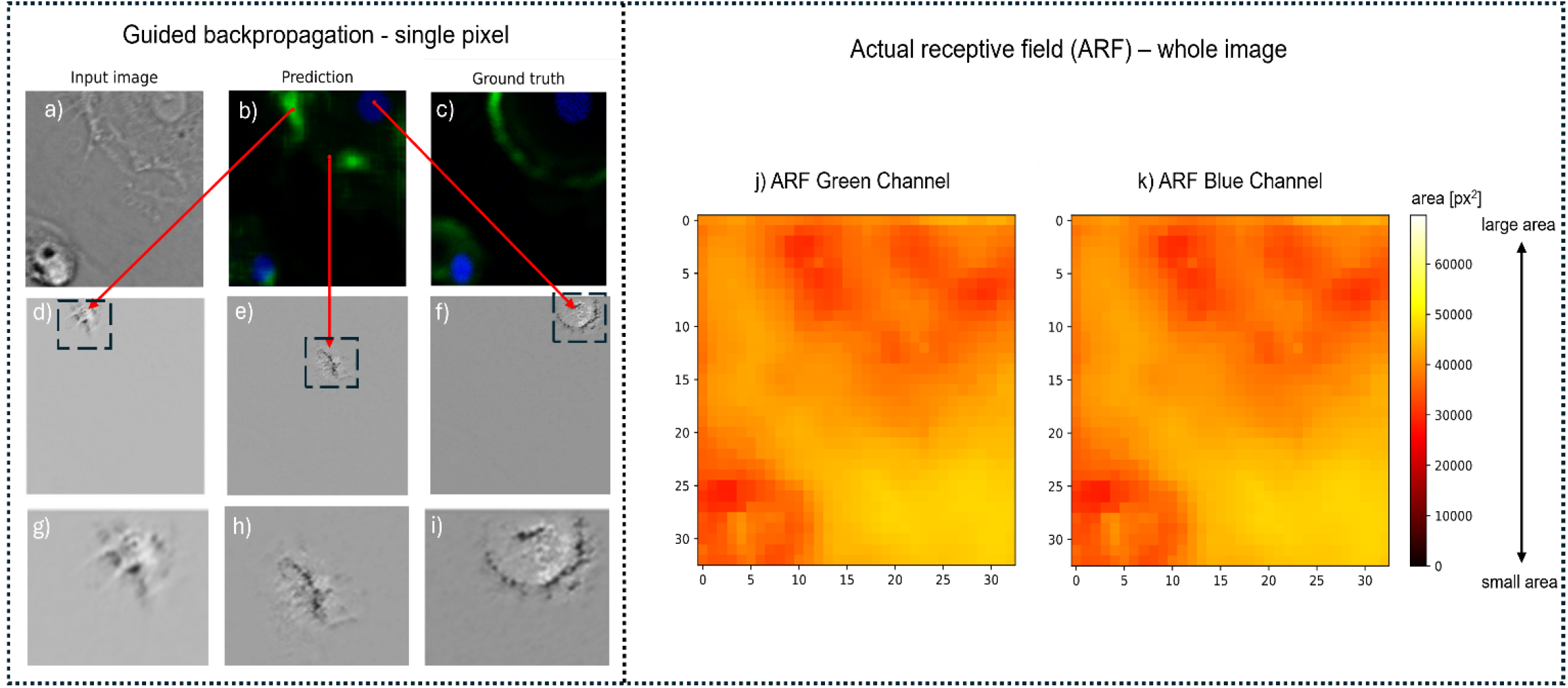
a)-f) Guided backpropagation (GBP) of the single pixels indicated in the prediction image b). The GBP of the blue colour channel (see f)) is part of a cell nucleus, revealing a circular receptive field, which contains relevant information for the staining decision. In contrast to this, the GBP of the cytoskeleton from the green colour channel leads to gradients having an irregular shape (see d) and e)). The GBP in d) origins from a correctly stained part of the skeleton, while e) is from a not recognised part, both GBS are calculated for the green colour channel. g)-i) provide zoomed in images of the marked regions in d)-f) showing details of the GBP results. In j) and k) the ARF sizes for every eight pixels in the image colour coded. The ARF for green and blue colour channel are nearly identical due to the shared network architecture for both colour channels.

A comprehensive analysis of all pixels within test images reveals that both blue and green channels typically require similarly large ARF to ensure accurate staining (Fig. 5 g) and h)). Here, the summarized ARF sizes were calculated for every eight pixels in the image, showing the in general smaller ARF for the cells and larger ARF for background regions. Most of the ARF areas are between 30.000 and 50.000 pixels, which show in general large ARFs for staining decisions. The largest areas are calculated for positions outside of cell structure. Unexpectedly, the ARF exceeds the TRF by multiples.

### Feature maps

To shed light into the discrepancy between ARF and TRF, we investigated the feature maps to better understand how spatial information is integrated across layers. Feature maps provide detailed insight into the activations of individual neurons during prediction. To analyse the network’s activation patterns in more detail, feature maps were extracted after each upsampling and downsampling block (see Fig. 2b). They reveal which image regions elicit a response in each block. In fig. 6 the degree of activation is coded by colours: the brighter a pixel (e. g. yellow), the greater the activation, while dark pixels indicate minor activation.

**Figure 6:**
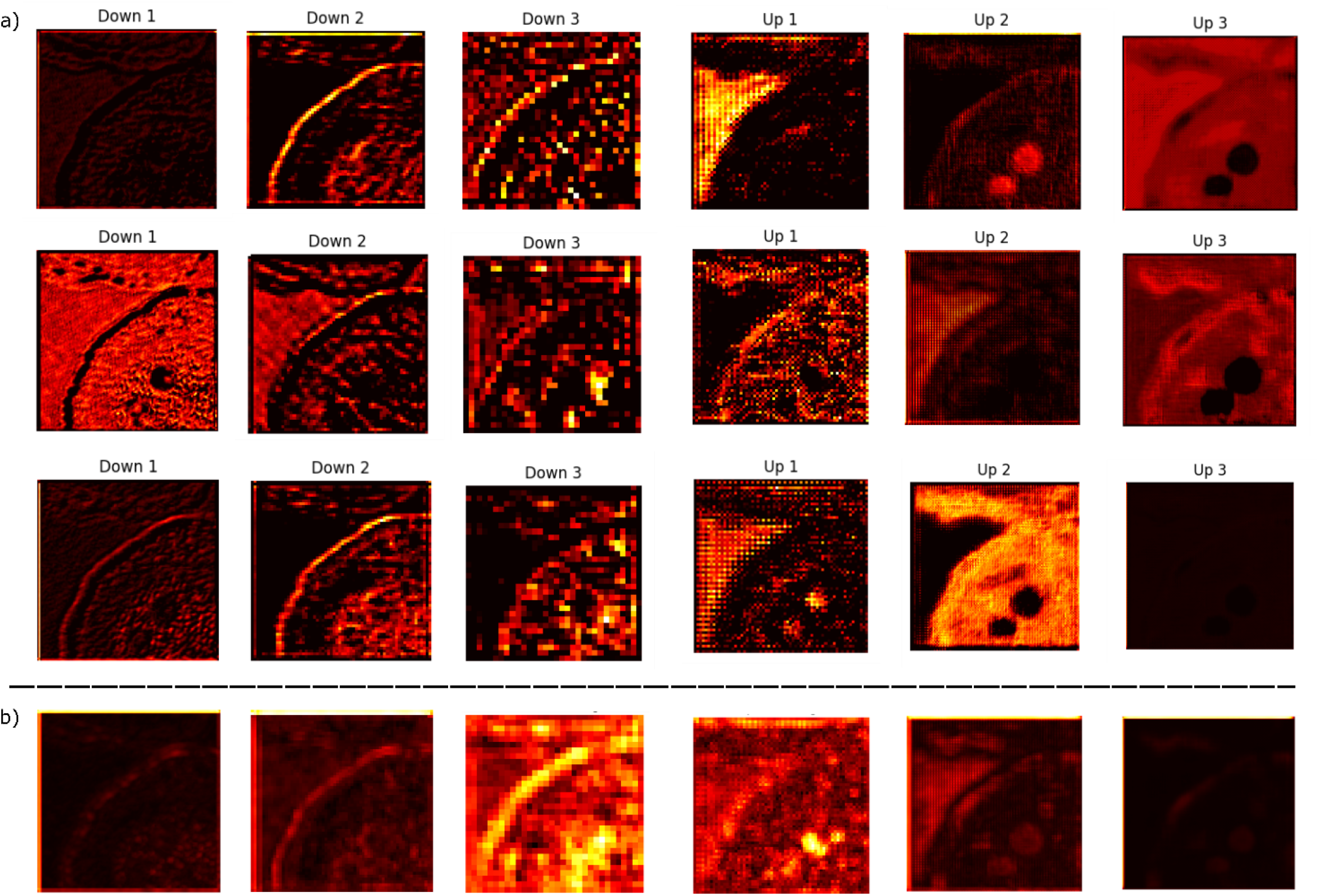
Feature maps provide insight in the activation in the network during inference mode; a) selected single channels and b) averaged feature maps over all channels after the indicated block. The resolution of the filtered images decreases with increasing network depth due to downsampling. After the third downsampling block, feature maps measure 32x32 pixels with 512 channels.

In early (Down 1) and late (Up 3) network blocks, minimal activation is observed in the averaged feature maps (see fig.6b)). Conversely, the highest activations occur in the deepest blocks (Down 3 and Up 1), where the features are highly compressed.

In the single channel feature maps, the spatial features of the cell such as the cell nuclei, cytoplasm and cell borders are shaping the activations (fig. 6a)). The cell nuclei are the most present in the late blocks (Up 2 and Up 3, layer details in fig. 2d)-e)), while the cytoskeleton, in particular the border of the cell is more prominent in the earlier blocks (Down 1 and Down 2, layer details in fig. 2c)). Unexpectedly, the background region, which does not contain pixels stained in the ground truth images raises significant activation. An example of this phenomena is visible in the block Up 1 in the first row of fig. 6a). It was much more expected that only the cellular shape and intensity variations from the cell organelles would cause activations.

Furthermore, the network is carrying out an internal segmentation although it was not trained for this specific task. In some maps the nuclei are segmented by activation while in others the cell border is activated. A spatial variance analysis revealed high variance in the maps after certain blocks. This is particular evident after the blocks Down 3 and Up 1, which are the deepest in the network and process highly compressed features (see fig. 7). The results of an additional principal component analysis with 3 components are provided in table 2 of the supplement. It reveals that the outermost blocks have components with the largest variance influence, while deep blocks such as Down 3 and Up 1 have components with minor influence. This result suggests that the outer blocks extract structure and patterns while the inner blocks have a high level of abstraction due to the previous compression.

**Figure 7:**
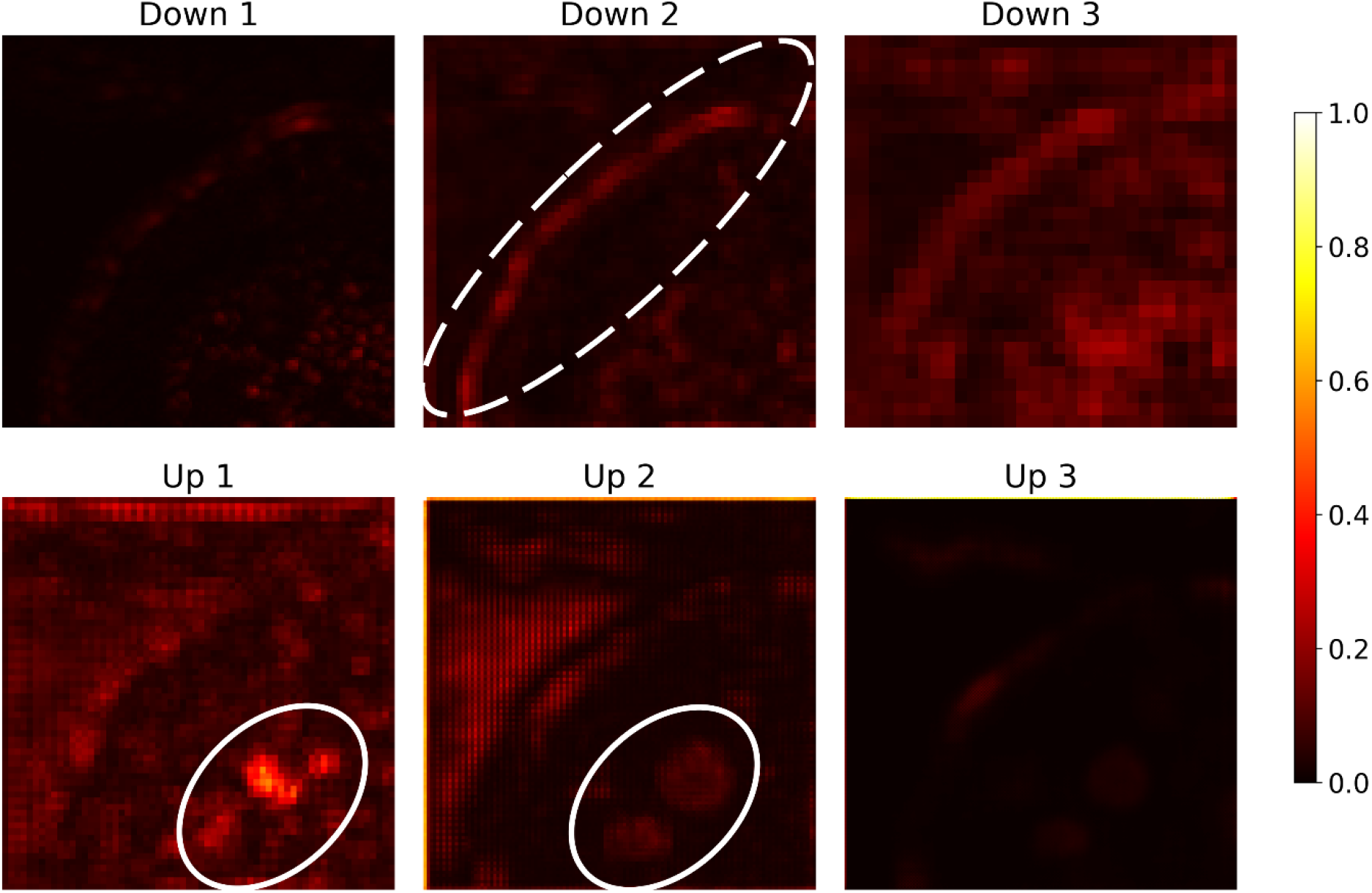
Spatial variance analysis of the feature maps after each block. Dashed white oval indicated recognized cell border and solid white ovals indicate recognized nucleus structures.

## Discussion

Our results show that the GAN-based model can generate biologically plausible virtual stains without the explicit need for segmentation. The explainability studies revealed that texture patterns of the cytoskeleton and background as well as the surrounding area of the nuclei strongly influenced the model’s predictions. This enables interpretation beyond pixel-wise similarity and supports its potential to replace selected chemical stains in standardized workflows. Using osteoclasts stained with Phalloidin and DAPI as ground truth, the network reproduced the green channel’s intensity distribution in the blue channel (Fig. 3g, h), likely due to shared weights across channels with a final RGB split in the last network block.

Next to the qualitative comparison of the predictions, the test metrics of SSIM>0.8 and PSNR>22dB support the finding, that the network can reconstruct the staining based on the intensity-only input images correctly. In future studies, phase information could be included to enhance the information density in the input modality. Reduction of the architecture complexity did negatively affect the test metrics, although the results are still comparable to the findings of other studies for virtual staining of H&E^20,21^.

In addition, we observed hallucinations of the trained network in the predictions on the test dataset, which are a known problem in deep neural networks^33^. These issues might be addressed by using a larger training dataset, unfortunately limited data is one of the main hurdles in training neuronal networks for biomedical tasks. Although there are some biomedical datasets accessible to the public, not enough staining, cell types and in particular not enough matched image pairs of stained and unstained cells are available. This issue can be investigated further using synthetic datasets, but this would introduce a reality gap between the training and test images, which is likely to lead to reduced performance of the network.

We address the explainability problem of neural networks by looking deeper into the trained network step by step. First, the predictions of the trained network based on augmented input images are tested, proving that simple structures such as rings and circles can leads to prediction of cells. So, this technique is a possible approach to create synthetic training datasets for different virtual staining applications.

Augmentation studies did find that the network learned to colour rings of 20 pixels radius, high intensity and 7 pixels border width strongest. Here, a clear maximum of the parameters in our study was found. In contrast, the circles with high frequency distribution of low intensity values, which were used as augmentations for the green coloured cytoskeleton did not lead to such obvious conclusions. This implies, that the imitation of the cytoskeleton in the input images is much more complex than the imitation of nuclei. A possible explanation for this finding could be the overall higher spatial distribution and complexity of the structures in the cytoplasm compared to the nuclei.

Followed by this initial study, which did only take forward processes in the network into account, we investigated backpropagation through the trained network. This led to the findings about ARF, describing important areas in the input image for the predictions made in the output image. The activations of whole layer blocks and single layers of the trained network are examined while processing an input image from the test set. All information from the input image is processed by the same layers and the colour channels are separated in the final upsampling block at the end of the network. This might explain the very similar ARF distributions and learned intensity probabilities. The given results emphasize that one network path per used colour channel should be trained. This could be realized using separate networks or integrating extra blocks into an existing architecture and thereby employing different layers and weights for different colour channels and stains. This concept was already demonstrated in biologically plausible spiking neural networks^34^ as well as pyramidal neural networks^35^ for image processing.

The analysis of ARF revealed that the predictions of cytoskeletal structures require the same size of receptive field compared to the prediction of the cell nuclei. The background part of the images needs more information and can only be reconstructed using a larger ARF. This was not expected, since the background of a stained image does not include any green or blue colour. We suggested that the intracellular compartments have a more complex structure and therefore need more information and larger ARF for a correct prediction. However, correct colouring of the background needs information about the position of cell borders, which requires a larger ARF than prediction of cell structures, which rely on less spatial information. This hypothesis needs further investigation and can be addressed by studies on a larger dataset and adapting gradient-based class activation mapping^36^. Furthermore, augmentations with higher complexity mimicking a whole cell with nucleus and organelles might help to determine the influence of input structures more realistic. Compared to training data for H&E staining, which is mainly used for tissue sections, fluorescent stained cell cultures have a larger portion of background in the whole image. This background has no signal in the ground truth images, so the fluorescent structures in ground truth are sparser than the structures in H&E ground truth images. Indeed, the large ARF for the background regions suggests, that these areas have large influence fields and need an unexpected high amount of surrounding information for correct staining prediction, this is supported by the internal segmentation revealed by the spatial variance analysis. These findings underly the important difference between intracellular versus extracellular image regions, which needs to be considered during training and for choosing test metrices. This study is based on mono-cell-cultures with only one layer and a large proportion of background, if the data becomes more complex, e.g. by adding a matrix material or using multi-cell-cultures, additional input information such as phase or polarization might further improve the training process^37^.

The averaged feature maps showed that the deepest network layers exhibited the most significant activation during inference. A possible explanation is the distillation of features during the downsampling. In the downsampling blocks, the features are not only extracted from the input, they are also compressed and analysed, resulting in less but more intense activated neurons in the layers of the third downsampling block. Interestingly, activation was also present in background regions of the image, which were not directly stained. This is consistent with the findings from the receptive field study and emphasizes the relevance of background and surrounding areas in the predictions. It suggests that the model integrates contextual information beyond the immediate stained structures, highlighting the complexity of the decision-making process. This contextual information is extracted very well based on the used input images, e.g. the cell border produces very specific activation patterns showing activation only at the cell border and no other region (see Down 2 in fig. 6b)), the same was observed for the nuclei and background. This leads to the conclusion that the intensity differences in the employed input images are sufficient for a successful training of networks. In general, the intensity contrast is crucial in the selection of training and testing data.

Even though brightfield imaging is limited in penetration depth and resolution compared to advanced label-free modalities such as SHG or two-photon microscopy, it is widely available and standardized in cell culture laboratories, making it a practical basis for virtual staining approaches. Our results indicate that such methods could reduce the need for chemical stains while remaining compatible with conventional antibody-based techniques, although current applicability is limited by the use of a single cell type and the occurrence of hallucinations in the generated stainings.

In the future, further studies on different cell types and other staining protocols may lead to networks, which can generalize better. So, multicell samples and multiple staining types can be covered by a single network. Additionally, combining VS with ultra-thin endoscopic technologies^38-41^ holds great potential for in vivo diagnosis. This integration could enable real-time, minimally invasive labelling of tissue using virtual staining, providing a powerful tool for clinical applications while minimizing the risk of infection, procedure related complications and the overall diagnosis time. In most biomedical applications large training datasets are not available, so transfer learning is a well-known strategy for data-reduction. In the case of virtual staining most models are trained for H&E staining and one approach for future work is to test whether these models can be additionally trained for IHC staining. Progress in the field of explainable neural networks is not only important for applications in biomedical research, it is also crucial for ethical considerations and safety issues in diagnostics assisted by artificial intelligence.

## Materials and methods

### Samples

Osteoclasts were differentiated from peripheral blood mononuclear cells which were isolated from human buffy coat of two different donors (purchased from German Red Cross, Dresden) by density gradient centrifugation as previously described ^42^. Cells were cultured in α Minimum Essential Medium (αMEM, Gibco) plus 2 % heat inactivated FCS and 100ng bone morphogenetic protein 2 (BMP2, Gibco) for 14 days in a triple culture with osteocytes and osteoblasts. A detailed protocol is described in ^43^. All cells were incubated in 5% CO2 and 95% humidity. The medium was changed twice per week. For detachment of cells, flasks were rinsed twice with phosphate-buffered saline solution (PBS, Gibco) before applying 0.25% trypsin/ 0.02% ethylenediaminetetraacetic acid (EDTA) solution (Gibco). Afterwards, cells were seeded in 50μL drops in glass slides (ibidi GmbH, Gräfelfing, Germany) for 1 hour to ensure adherence before adding medium and culturing for 14 days.

### Fluorescence microscopy

For every staining, after washing with PBS, cells were fixed with 4% formaldehyde in PBS (SAV LP GmbH, Flintsbach a. Inn, Germany) for 30 minutes and washed with PBS. For permeabilization, 0.1% Triton X-100 in PBS was applied for 5 minutes and samples were washed five times with PBS, followed by blocking with 3% Bovine Serum Albumin (BSA) (Carl Roth, Karlsruhe, Germany) in PBS for 1 hour. Then cells were incubated with Phalloidin iFLuor 488 (1:1000, Abcam, Cambridge, UK) and DAPI (1 μg/mL) (AppliChem, Darmstadt, Germany) for 1 hour protected from light and subsequently washed with PBS. Samples were washed again with PBS and finally imaged using a confocal laser scanning microscope (STELLARIS 8 from Leica Microsystems GmbH, Wetzlar, Germany). For all image pairs in the datasets, each imaged location was acquired in three channels: blue, green and brightfield each with a resolution of 4096 x 4096 px. Several spots in multiple wells of the ibidi slides were imaged, with seven spots acquired for each donor. The signal intensity was adjusted to compensate for inherent staining variability, specifically by balancing over- and under-illumination.

### Image processing

The acquired images were cut down into 256 pixels by 256 pixels images and split into 4515 image pairs for training and 210 image pairs for testing. The training set was further split into 70% training images and 30% validation images. During training and testing all images were normalized before feeding into the networks. The blue colour channel of the stained images was smoothed using a Gaussian filter with a standard deviation of two pixels. Input images are 1-channel grayscale images, while predicted output images are 3-channel RGB-images of the same height and width.

### Network training

A GAN was trained for 150 epochs based on the dataset described in the methods section. The dataset consists of widefield and fluorescence images of osteoclast cultures. In the ground truth images, the cells are stained using DAPI and phalloidin.

As generator network in the GAN architecture a Pix2Pix architecture^44^ with three up blocks and three down blocks was used, before merging the skip connections to the up blocks, an attention block was added ^45^ (see fig. 2). The learning rate of the generator was set to 10^−4^ and kept constant using an Adam optimizer. The generator network takes an unlabeled widefield image as input and translates it into a stained image. The generator loss was defined according to eq. (1), here *MSE*(*y*_*gt*_, *y*_*pred*_) is the mean squared error between the ground truth image *y*_*gt*_ and the predicted image *y*_*pred*_. *D*(*x*, *y*_*pred*_) represents the resulting probability of the discriminator for a fake image, produced as prediction from the generator network to be a real image. The metrics in validation and testing were defined as SSIM and PSNR.

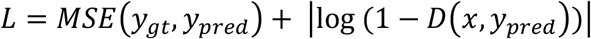

For the discriminator a PatchGAN architecture was employed. As the final layer a sigmoid activation layer was chosen. It was trained using a constant learning rate of 10^−5^ and an Adam optimizer. The dataset was fed into the networks with a batchsize of 32 images and the training was performed using a NVIDIA RTX A6000 GPU.

In fig. 8 a final result of a large test image is presented. Although the prediction of cytoskeleton (colored in green) shows issues, the prediction of nuclei is realistic from a quantitative point of view. The corresponding input images as well as the full prediction and ground truth images can be found in fig. 1 of the supplement.

**Figure 8:**
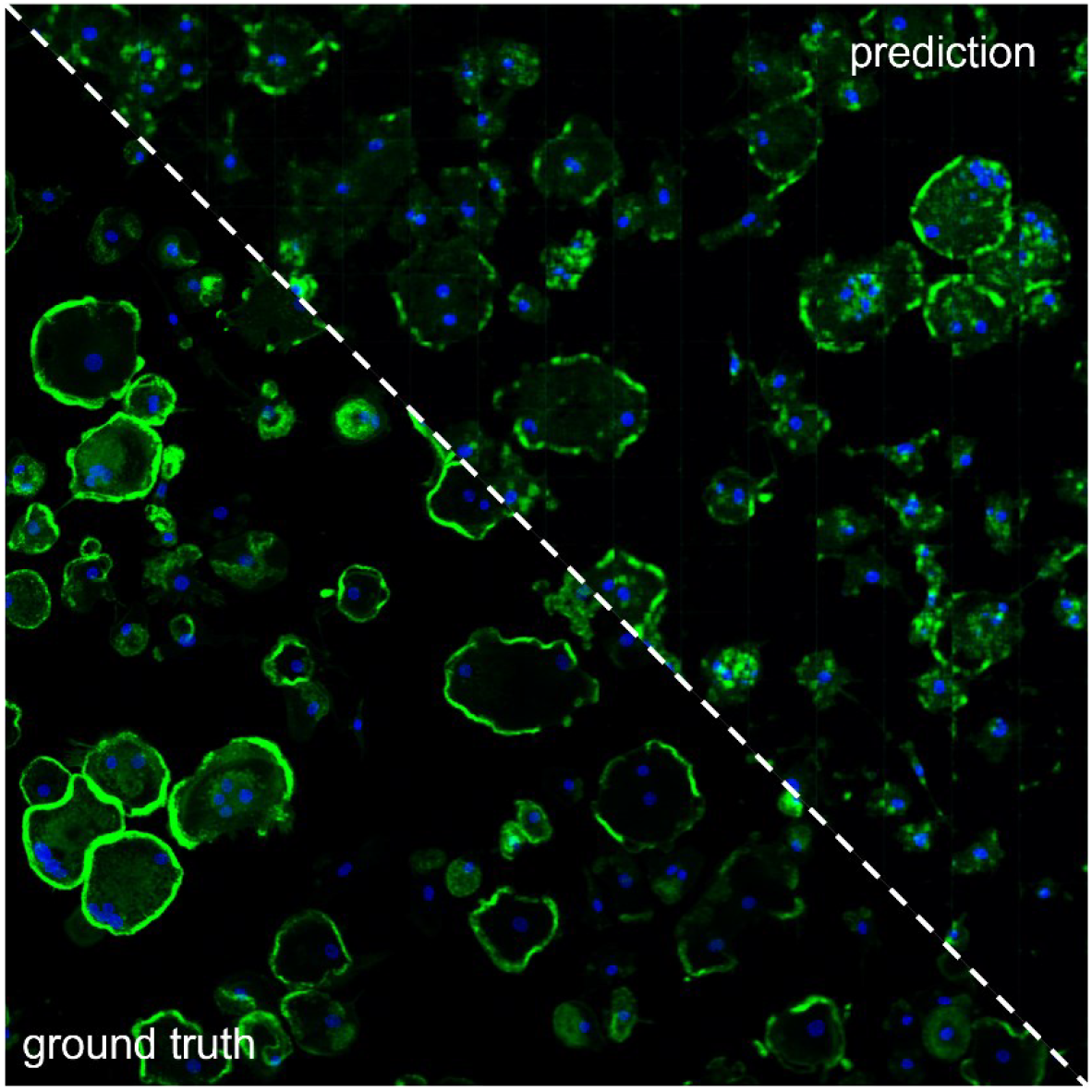
Prediction and ground truth of a large 4096x4096 px image. For the prediction, it was cut into 256x256 px snippets and afterwards stitched together.

### Augmented input images

Augmented structures were added to input images in order to determine which structural image components of the input image are coloured blue or green in the network prediction and thus recognized as cell components by the network.

The cell nuclei appear as dark rings in the input images of the training and test data set, so rings of different sizes, intensities and with varying border width were used as augmentations. The cell body and cytoskeleton are imitated as noisy regions and varied in size and intensity. These augmentations were added to an input image using the image processing toolbox from The MathWorks Inc.^46^. Examples of augmented input images and their corresponding predictions are shown in fig. 9.

The resulting predictions made based on images with the above-described augmentations are evaluated by calculating the mean intensity of a certain colour channel over the region of the added structure.

## Acknowledgements

The authors thank David Krause from the Chair of Measurement and Sensor System Technique (TU Dresden) for assistance in the TRF calculations and Dmitry Richter from Wellmann Center for Photomedicine (Massachusetts General Hospital) for the discussions about the network training in general. Furthermore, the authors recognize the support from DFG (project CZ55/54-1), Development bank of Saxony (application 100589210) and the German Federal Ministry of Education and Research (BMBF) for funding of the initiative ProMatLeben and particularly the subproject “Innovative polymer composites for patient individual additive Manufacturing of biodegradable dental implants (InnoPoly)” (grant number 13XP5067).

Fig.1 was created in BioRender: Schmidt, K. (2025) https://BioRender.com/3acbmz9.

## Author contributions

M.v.W. prepared and imaged the samples; K.S. and A.O. trained the network and implemented the code; K.S., N.K. and M.v.W. analysed and discussed the results; K.S. and M.v.W. prepared the manuscript; all authors revised the manuscript; M.G. and J.C. supervised the project;

## Data and Code availability

The neural network model as well as the evaluation code is available at https://gitlab.hrz.tu-chemnitz.de/anob943c-at-tu-dresden.de/ganvis-public. The training and testing dataset is can be downloaded from figshare (DOI 10.6084/m9.figshare.29625476).

