## Supplement for "Understanding Virtual Staining with generative adversarial networks for Osteoclast Imaging"

### Supplement to the research paper „Understanding Virtual Staining with generative adversarial networks for Osteoclast Imaging"

#### 1. Prediction of large image

A large grayscale image (4096 x 4096 px) was used as input for the trained network. For processing, the image was cut into 256 x 256 px snippets and each snippet was taken for a single prediction. Afterwards the predicted images with the same size as the input image, but 3 color channels were stitched together again to create the complete prediction of a large image (shown in fig.1).

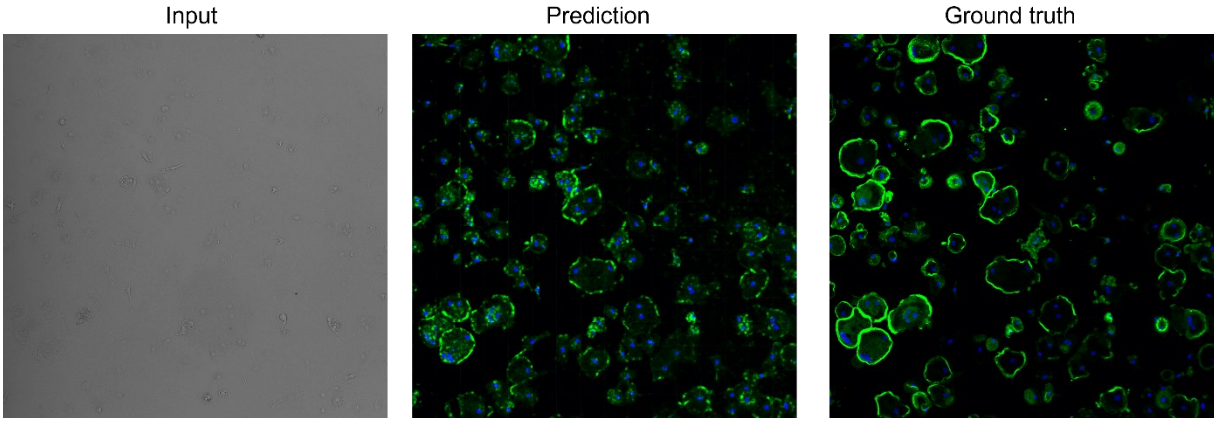


*Figure 1: Prediction of a 4096x4096 px image.*

#### 2. Zoom-Ins and hallucinations

Zoom-Ins from the test images in fig.3 of the manuscript and corresponding highlights of hallucinations (red ovals in the zoomed regions) are provided in fig.2.

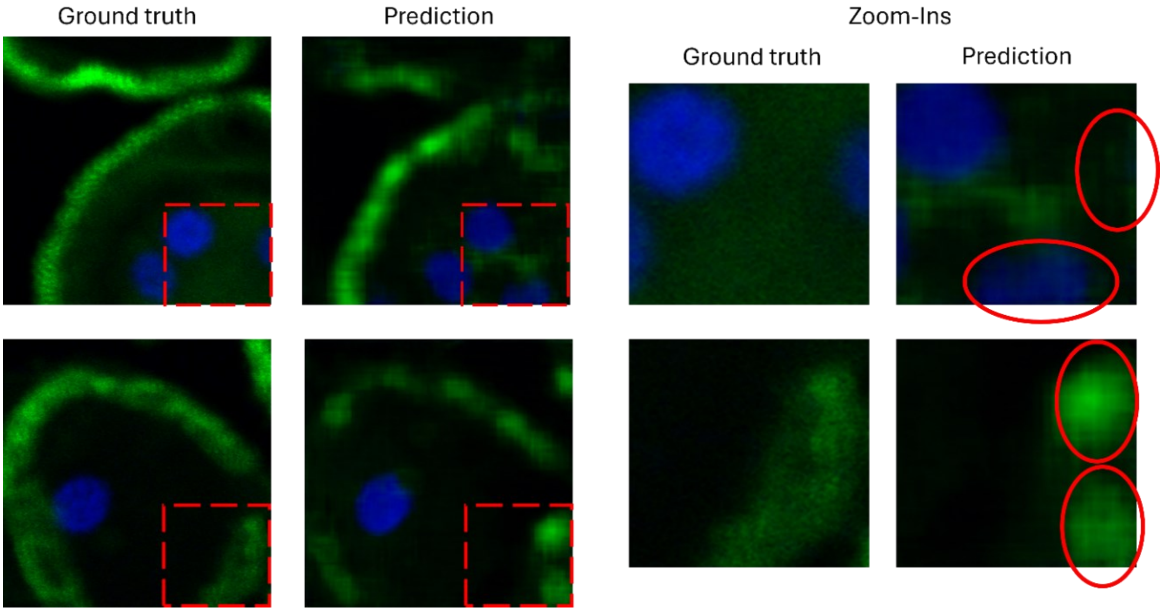


*Figure 2: Zoom-in from hallucination areas in the predictions of the test dataset. Ovals indicate regions with hallucinations.*

#### 3. Theoretical receptive field size

The method for calculation of the theoretical receptive field (TRF) is based on the guide by Dumolin and Visin [R1]. Here, the output size $o$ of an input image with size $i$ for a convolutional layer with kernel size $k$, padding $p$ and stride $s$ is defined as follows:

$$o = \left\lfloor \frac{i + 2p - k}{s} \right\rfloor + 1 \qquad \text{Eq.1}$$

To derive the TRF, backpropagation from output plane to image plane instead of forwarding from input to output plane is necessary. Due to this requirement, the padding is neglected, since backpropagation from a pixel in the middle of the image is assumed to derive the maximum TRF size. This simplifies eq.1 to:

$$o = \left\lfloor \frac{i - k}{s} \right\rfloor + 1 \qquad \text{Eq.2}$$

The Pooling layers in the Down blocks of the Pix2Pix generator perform a maximum pooling of the input plane to the output plane and thereby reduce the size. This can be expressed by eq.2 as well, if padding is neglected. According to the appendix of the publication from Loos et. al [R2], activation functions and attention gates have no influence on the TRF.

For the calculation of TRF it is assumed, that due to deconvolution processed in the upsampling blocks of the architecture, only a single pixel in the feature maps between downsampling block 3 and upsampling block 1 is relevant. All relevant layers, kernel sizes, stride sizes and plane sizes are given in tab.1.

Based on the calculations in tab.1, the TRF is 96 pixels, which results in an TRF area of $96^2 = 9216\,px$.

*Table 1: Relevant layers for TRF calculation, their properties and results of input size.*

| Block | Layer | Kernel size | Stride | Output size | Input size |
|---|---|---|---|---|---|
| Down 1 | Convolution2d | 4 | 1 | 1 | 4 |
| | Convolution2d | 4 | 1 | 4 | 7 |
| | MaxPooling | 2 | 2 | 7 | 14 |
| Down 2 | Convolution2d | 4 | 1 | 14 | 17 |
| | Convolution2d | 4 | 1 | 17 | 20 |
| | MaxPooling | 2 | 2 | 20 | 40 |
| Down 3 | Convolution2d | 4 | 1 | 40 | 43 |
| | Convolution2d | 4 | 1 | 43 | 46 |
| | MaxPooling | 2 | 2 | 46 | 92 |
| Inc | Convolution2d | 3 | 1 | 92 | 94 |
| Inc | Convolution2d | 3 | 1 | 94 | 96 |

#### 4. Actual receptive field calculation

To gain insights into the activation and experimental influence of input pixels on the predicted values in the RGB-output, guided backpropagation was used [R3]. By this, the influence gradients of one output pixel were backtracked to the input plane. An example calculation of gradients for an output pixel in the middle of black circle is presented in fig. 3a. The scale represents the intensity of gradients. From this figure is visible, that the largest gradients are located around the position of the used output pixel.

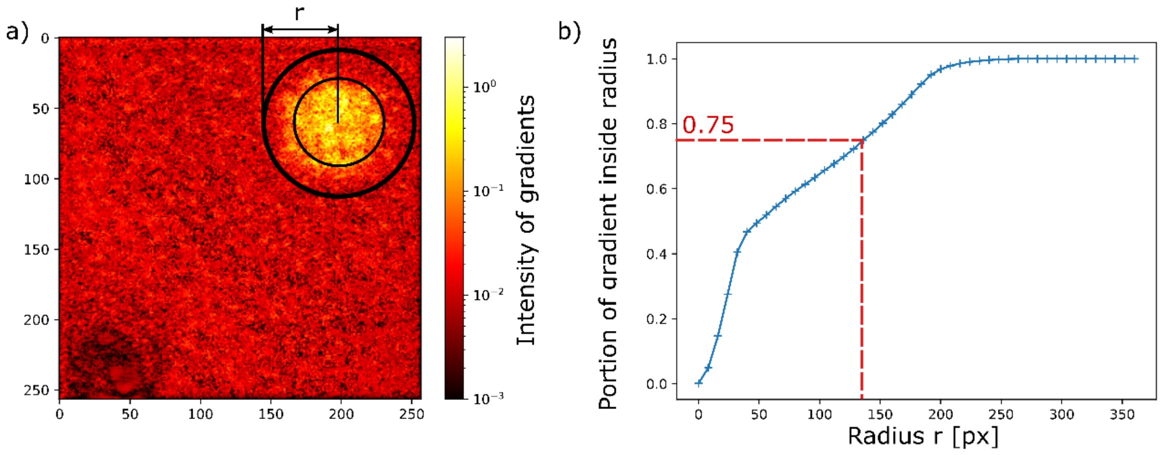


*Figure 3: actual receptive field calculation*

The radius r of the actual receptive field (ARF) size is calculated based on the portion of gradients inside the area around the relevant output pixel. This is demonstrated by circles with different radii in fig. 3a, where different portions of the gradients are inside circles of different radii. In fig. 3b, the link between portion of gradient and resulting radius of ARF is shown. The radius of the area including 75% of the relevant gradients is about 135 pixels (refer to red dashed line in fig. 3b).

#### 5. Principal Component Analysis

A principal component analysis for the feature maps extracted after each block in the pix2pix architecture reveals large variance components for the outer blocks and smaller components for deeper/inner blocks.

*Table 2: Results of the principal component analyses with 3 components.*

| Block | PC1 (%) | PC2 (%) | PC3 (%) |
|---|---|---|---|
| Down 1 | 44.44 | 15.39 | 9.66 |
| Down 2 | 36.23 | 17.71 | 5.38 |
| Down 3 | 6.36 | 5.49 | 5.46 |
| Up 1 | 11.35 | 6.34 | 5.85 |
| Up 2 | 27.51 | 14.98 | 11.30 |
| Up 3 | 45.24 | 19.98 | 13.39 |
